# Visualization of the architecture of flexible chromatin and the binding of chromatin regulators

**DOI:** 10.64898/2026.06.05.730377

**Authors:** Haonan Zhang, Yan Li, Chengcai Pan, Fu Bo, Cong Yu, Weiwei Niu, Huimin Yang, Kai Song, Ping Zhu

## Abstract

Chromatin organization plays a central role in regulating genome accessibility and gene expression in eukaryotic cells. However, the inherent flexibility and structural heterogeneity of chromatin pose significant challenges for its structure determination. Here, we use a Nuc-back strategy with cryo-electron tomography (cryo-ET) and subtomogram averaging methods to visualize chromatin at the nucleosome level by averaging nucleosome at moderate-to-high resolution, classifying the fundamental unit of chromatin, i.e., nucleosome, into distinct classes, and linking different nucleosome structures to chromatin architecture. We reveal that nucleosome heterogeneity is a key factor in chromatin flexibility which disrupts interactions between nucleosomes. In addition, this strategy allows for the localization and visualization of chromatin regulators and their structure on chromatin. These results provide a foundation for future research in 3D genome and epigenetic process visualization.

**Graphical Abstract:** 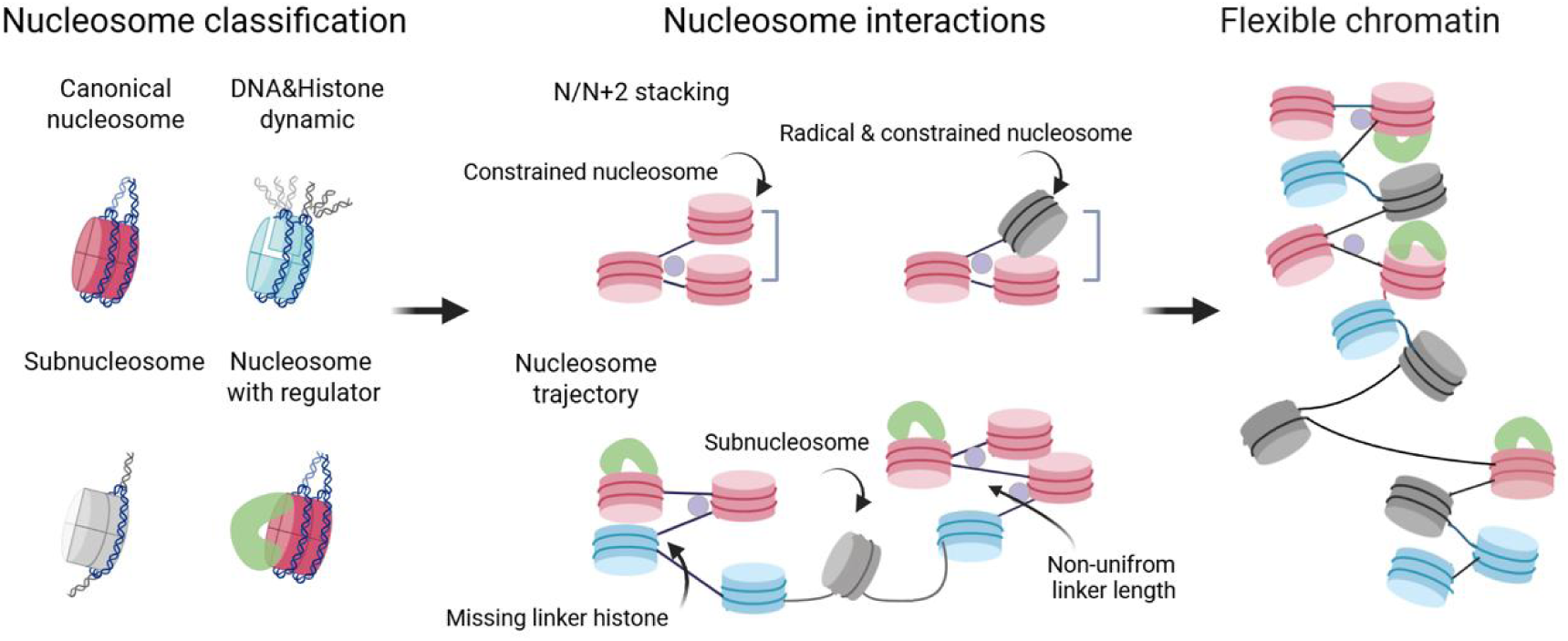

## Introduction

In eukaryotic cells, nucleosome serves as the fundamental repeating units of the chromatin. Each nucleosome consists of a nucleosome core particle (NCP), where 147 base pairs (bp) of DNA are wrapped around an octamer of histone proteins (comprising two copies each of H2A, H2B, H3, and H4) forming approximately 1.7 left-handed super helical turns [1]. These nucleosome cores are connected by linker DNA, typically ranging from 10 to 90 bp, creating a “beads-on-a-string” arrangement with an 11-nm diameter [1]. Facilitated by linker histones and other factors, e.g., divalent cation or chromatin architecture proteins, the nucleosome arrays are further assembled to condensed chromatin, which typically show as fibers around 30-nm with a local two-start zig-zag arrangement of nucleosomes [2].

Recent advances in cryo-electron microscopy (cryo-EM) single-particle analysis (SPA) have provided high-resolution structures of NCP in complex with various factors [3–7]. These studies revealed that the overall NCP structure is relatively conserved across different DNA sequences. However, it has been demonstrated that nucleosomes also possess intrinsic structural plasticity, characterized by rapid movements of DNA and histone regions, a phenomenon known as “nucleosome dynamics” [8,9]. These dynamics alter the linker DNA length and conformation through breathing, splitting, and sliding motion of the nucleosome DNA thus inducing the chromatin plasticity. Furthermore, nucleosomes *in vivo* exhibit significant compositional diversity through post-translational modifications or histone variant exchanges [10,11]. These variations of local charge or conformation not only alter the nucleosome structure, but also have impact on the interactions between nucleosomes and its binding proteins. Therefore, in analyzing chromatin structure, nucleosomes should not be treated as a single homogeneous class.

Constructed from dynamic nucleosomes, poly-nucleosomes or chromatin fibers are typically flexible as they present the interplay among various nucleosomes with changing structures. For *in vitro* studies, we have determined the structure of chromatin particles reconstituted with 12× nucleosomes and linker histones H1 or H5 at high resolution using single particle analysis [12,13], which employs defined nucleosomes with 601 DNA sequence, with accurate linker histone/nucleosome ratios, and computational classification yielding homogeneous particles. For the structural study of longer chromatin fibers, cryo-electron tomography (cryo-ET) combined with sub-tomogram averaging (STA) has emerged as a powerful approach. Our previous study on reconstituted chromatin fiber with 50x nucleosomes *in vitro* revealed a regular subunit of 9 nucleosomes through STA [14]. However, the determination of poly-nucleosomes structure is limited by the particle homogeneity and involves high computational complexity, particularly due to the need to account for flexibility between multiple nucleosomes, many particles are often excluded from averaging. Recently, the study using cryo-ET to investigate individual nucleosomes within chromatin condensates has revealed how linker length regulates condensate behavior, providing a promising research approach for studying chromatin *in vitro* [15,16].

For *in vivo* chromatin, whose composition is heterogeneous, exhibits much more irregular nucleosome arrangement, even at single nucleosome level. Studies on purified mitotic chromosomes using cryo-EM revealed that linker DNA length and entry/exit angles vary broadly, suggesting that the nucleosome particle represents only an average state rather than a uniform structure [17]. Moreover, cryo-ET combined with denoising uncovered nucleosome stacking is highly heterogeneous and dynamically regulated, for instance by Mg²⁺ concentration or PAD4-dependent citrullination [18]. Cryo-focused ion beam (Cryo-FIB) milling is predominantly adopted to study macromolecular complexes in their native environment [19]. In HeLa cells, STA mapped back onto raw tomograms show that nucleosomes in heterochromatin follow irregular paths [20], whereas in yeast, extracted chromatin can display canonical nucleosome structures, but *in situ* nucleosomes often deviate from the classical conformation, highlighting the dynamic and environment-dependent nature of nucleosome organization [21]. In RPE-1 cells, ordered dinucleosome stacks are observed, but higher-order stacks are heterogeneous [22]. Studies of human T lymphocyte chromatin fibers further show that nucleosomes (with or without H1), form typical structures within relaxed, variable zig-zag organizations [23]. Together, these findings emphasize that native nucleosome positioning and interactions are variable and plastic, contributing to highly irregular chromatin architecture.

Many studies have employed a strategy of averaging nucleosomes followed by mapbacking to observe chromatin; however, such studies often remain at a macroscopic level and fail to establish a direct link between nucleosomes and chromatin architecture. Moreover, the visualization of nucleosome *in vivo* at high resolution is proved quite challenging. The low signal-to-noise ratio (SNR), caused by the relatively small molecular weight of nucleosome (∼250 kDa) and the noisy nuclear environment, makes it difficult to determine the accurate spatial parameters, such as nucleosome coordinates and orientations, in chromatin context.

In addition, the binding of various epigenetic regulating factors to the chromatin plays a key role in regulating gene expression. The structures of nucleosomes in complex with different protein factors have been extensively studied *in vitro* by SPA and many of them have been resolved at near-atomic resolution [3]. However, locating these regulating factors within the chromatin context is methodologically challenging, particularly for low-abundance factors. Moreover, the binding modes of protein factors to the *in vivo* chromatins and the consequent chromatin structural changes at their binding site are much less understood.

In this study, we applied cryo-ET combined with STA to analyze the type, location and orientation of individual nucleosomes within chromatin. These analyses were performed using chromatin reconstituted *in vitro*, chromatin extracted form nuclei, and *in situ* cryo-lamellae, thereby demonstrating the potential to resolve nucleosome structures at moderate-to-high resolution in the chromatin context by employing extensive classification to distinguish diverse categories of nucleosomes and reveal their dynamics. Based on this, we introduced a Nuc-back strategy, which achieves a comprehensive chromatin map by localizing and mapping the nucleosome category motifs back into the reconstructed chromatin volume. By applying the Nuc-back strategy from *in vitro* to *in vivo*, it enables us to link nucleosome categories with chromatin architecture. We demonstrated that nucleosome heterogeneity is an important factor in chromatin flexibility through disrupting interactions between nucleosomes. As an extension, we further applied this strategy to visualize how protein factors bind to chromatin and subsequently alter the local structure at their binding site. These findings enhance the understanding of chromatin organization at the single-nucleosome level and provide a foundation for future studies on epigenetic regulation process.

## Results

### Determining accurate spatial parameters of nucleosomes by STA with deep classification

To acquire accurate spatial parameters of nucleosome in the context of chromatin, we first reconstituted chromatin fibers *in vitro* as a reference to optimize the STA and classification procedure. Chromatin particles with 12×177 bp nucleosome repeat length (NRL), compacted by linker histone H1 (i.e., 12×177 bp chromatin particle, dataset 1), were prepared following the protocol as previously described [12]. The samples were vitrified and subjected to cryo-ET data collection, and nucleosome densities within the chromatin context were extracted for STA analysis. A total of 108 tomograms were collected (see Table S1). Multi-round classification and iterative refinement in M [24] and Relion [25] yielded a final resolution of 4.1 Å for NCP (Figures 1A and S1). The local resolution within the histone regions reached 3-4 Å, enabling visualization of side-chain features (Figure S2). We then attempted to resolve the linker histone density associated with the NCP. No-alignment 3D classification using a focused mask on the H1 region, while retaining a small portion of the core particle density, was performed. By adjusting the T-value and mask diameter, a distinct H1 density was identified at 4-5 Å resolution, which enabled visualization of its backbone structure (Figure 1A).

**Figure 1.**
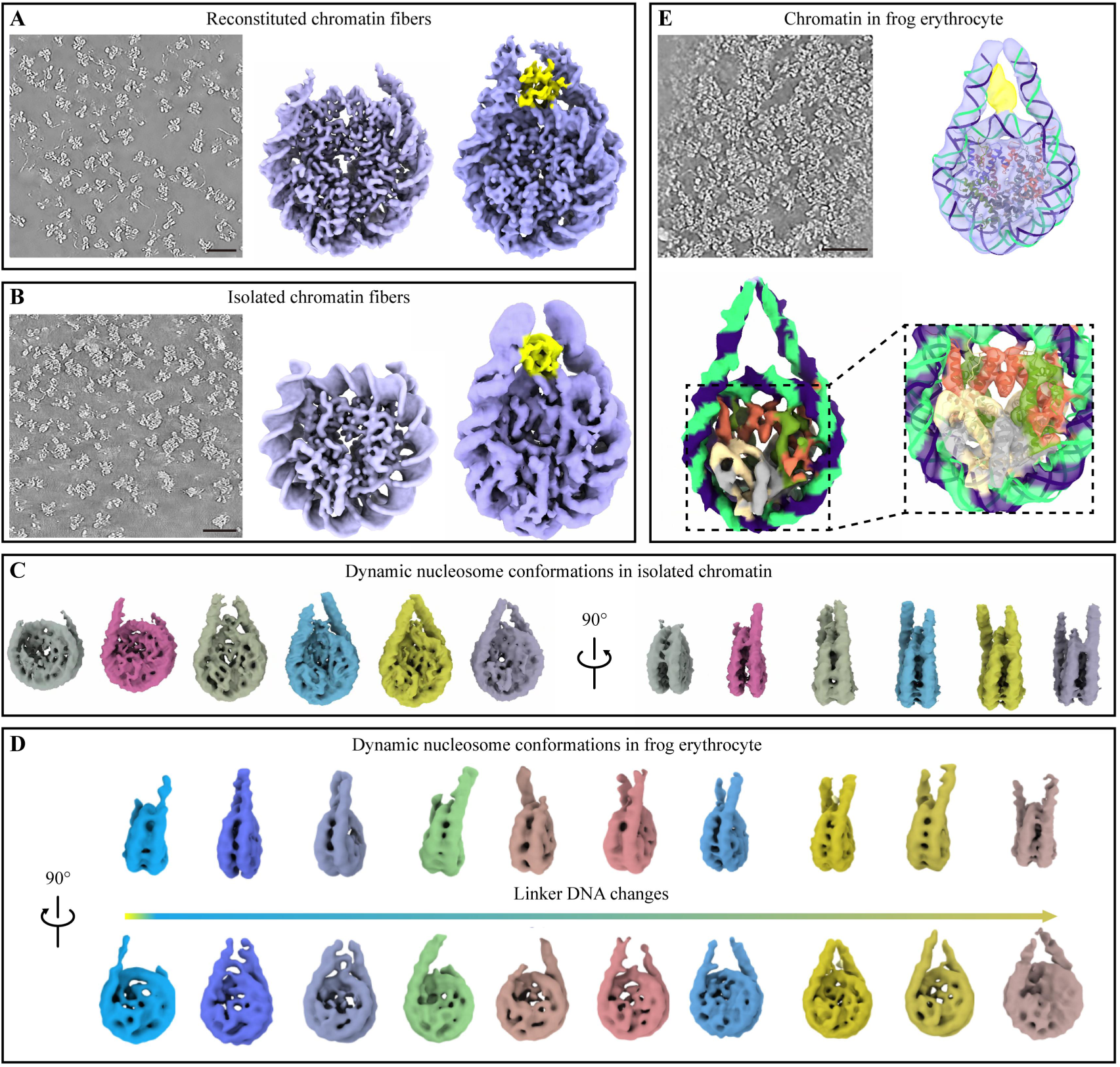
STA analysis of individual nucleosomes in chromatin from different sample sources. **(A-B)** STA results of nucleosomes in reconstituted chromatin (A) and isolated chromatin (B). From left to right, shown are a tomographic slice of chromatin, the averaged result of nucleosome core, and the averaged result of nucleosome associated with linker histone H1 (yellow), respectively. **(C-D)** Different nucleosome classes from the isolated chromatin (C) and chromatin in frog erthrocyte (D). **(E)** STA results of *in vivo* nucleosomes in the lamella chromatin from frog erthrocyte. Panels shown are the tomographic slice of chromatin, the averaged density of nucleosome associated with linker histone H1 (yellow) (up-right), the density of averaged nucleosome core particle (bottom-left) and an enlarged view fitted with the atomic model (PDB: 7DBP) (bottom-right). Scale bars: 50 nm.

Next, we applied a similar workflow to chromatin fragments extracted from SF9 cells which contain a more heterogeneous nucleosome composition (dataset 2). The 3D reconstruction revealed the extracted chromatin fibers of varying lengths are largely compact (Figure 1B). Using the STA workflow, we obtained a 5 Å resolution for the nucleosome core and a 6.4 Å resolution for the nucleosome containing H1 (Figures 1B and S2). At this resolution, secondary structure features in histones and the distinct double helix characteristics of the wrapped DNA are clearly visible. Interestingly, compared to the reconstituted chromatin, more nucleosome categories with distinct variations in linker DNA trajectory and histone densities were identified in the extracted chromatin (Figures 1C and S3), suggesting the extracted chromatin is more heterogeneous and with more dynamic compositions.

Beyond reconstituted and extracted chromatin, we further studied the structure of *in vivo* chromatin within the lamellae milled by cryo-FIB (dataset 3). Previous studies have shown that ice thickness significantly impacts the SNR and data quality, posing substantial challenges for analyzing small molecules such as nucleosomes at high resolution [26]. By testing lamellae thickness ranging from 70 to 220 nm, we found that the thickness of 70-150 nm provided sufficient SNR for resolving nucleosome features after extensive classification (Figure S4). We therefore analyzed *in vivo* nucleosome structures in frog erythrocyte nuclei using lamella within this thickness range. Non-nucleosome particles were discarded by manually defined boundaries, template matching, and multiple rounds of 2D and 3D classification (Figure S5). After 3D classification, ten distinct nucleosome categories were identified (Figure 1D). These categories differed primarily in linker DNA trajectory, showing variations in lengths and entry/exit angles, possibly reflecting intrinsic linker DNA dynamics or the influence of neighboring nucleosome interactions. Moreover, the histone octamer also fails to adopt a stable conformation. These results demonstrated that distinct nucleosome categories *in vivo* can be well distinguished, even within compact chromatin, by combining thin lamellae, STA and deep classification.

To improve the *in vivo* nucleosome structure features, we selected particles from one category displaying histone-like features and performed further refinement using the similar STA and classification process. The refinement yielded an average resolution of 9.5 Å, which enabling definition of histone secondary structures and clear distinction between histone and DNA densities (Figure 1E). An H1-bound nucleosome structure was also resolved (Figure 1D), although at a lower resolution (13 Å), likely due to the dynamic binding manner of H1.

Collectively, our results indicate that the location and orientation of nucleosome within chromatin, both *in vitro* and *in vivo*, can be determined by STA and extensive classification with sufficient accuracy to generate averaged nucleosome maps containing characteristic structural features. This lays the foundation for accurate nucleosome mapping-back (Nuc-Back) procedures and facilitates further chromatin analysis at the single nucleosome level.

### A Nuc-back strategy enables the visualization of irregular nucleosome arrangements

Having distinguished various nucleosome categories at medium to high resolution via STA, we obtained the metadata for each nucleosome. Next, we treat the averaged nucleosome densities in different categories as fundamental building motifs, which enables us to assemble the overall architecture of chromatin by putting the class-average density of the corresponding category back into the 3D space, referring to the orientation of each nucleosome obtained in the STA process. Subsequently, combined with the denoised tomograms, the orientation and positioning of nucleosomes are validated. Through this way, which we refer as Nuc-back strategy, the whole map of chromatin can be built.

Firstly, we tried to optimize the Nuc-back strategy using cryo-ET data of *in vitro* assembled tetranucleosomes (dataset 4) (Figures 2A and 2B). By classification, we defined three categories of nucleosome, including two categories in which the linker DNA densities were resolved on both sides (red) and on only one side (blue) respectively, and one category of poor resolved density (gray) (Figure 2D). Mapping the building motif, i.e., the averaged density of classified category, back to the tomographic volume using the corresponding center coordinate and orientation parameters from STA, we found they aligned very well with the raw densities (Figure 2E). Since the true orientation of nucleosomes is difficult to obtain (i.e., lacks the ground truth), we validate the accuracy of each nucleosome in chromatin by overlaying the denoised slices and volumes which shows an overall good consistency (Figures 2E-F). However, in spite that most nucleosomes were fitted and assigned to one of the categories, some nucleosome-like densities remained unassigned to any category. We designated these as missing nucleosomes (Figure 2E).

**Figure 2.**
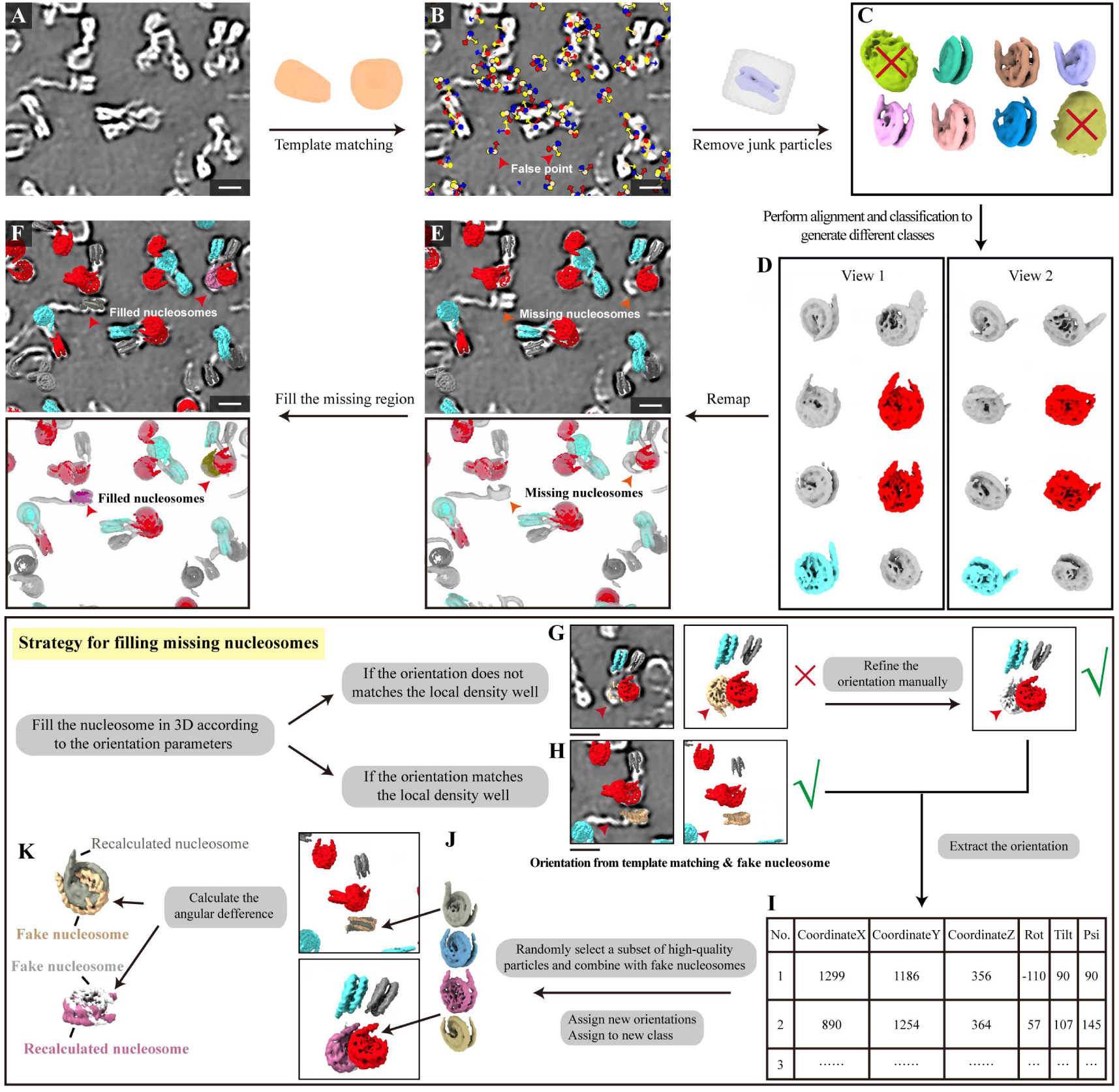
Workflow of the Nuc-back strategy. **(A)** Reconstructed tomogram. **(B)** Template matching to determine the initial coordinates and orientations of nucleosomes. **(C)** 3D classification to remove junk particles. **(D)** Fine classification of all nucleosome particles. **(E)** Mapping the classified category density map back onto the tomographic volume using the computed orientations, revealing some missing nucleosomes. **(F)** After filling the missing nucleosomes. **(G-K)** Strategy for filling missing nucleosomes. Visually assess the nucleosome orientations from template matching in the denoised tomographic slice, e.g., incorrect (G) or correct (H). If the orientation is incorrect, manual refinement will be involved according to the denoised tomogram. (I) Record the missing nucleosome orientations from (G) and (H). (J) Fill in the nucleosomes into the 3D space according to the newly generated classes. (K) Calculate the angle difference between the recalculated nucleosome orientations and the ‘fake’ nucleosome orientations. If the difference is less than 22.2°, the recomputed orientation was used to fill the missing nucleosome, otherwise the ‘fake’ orientation would be used. Scale bars:10 nm.

These missing nucleosomes were then subjected to re-identification by template matching or manually orientation refinement based on the denoised slice or segmentation (Figures 2G-K). Finally, iterative classification and orientation assessment were performed to ensure that all nucleosomes are properly located and mapped back. For detailed procedures, see Materials and methods.

By remapping all nucleosomes back to the tomogram densities, the overall architecture of the oligonucleosomes could be visualized, which are arranged in a closed zig-zag pattern and packed as a tetranucleosome unit (Figures 3A and S6A). Notably, the compositional heterogeneity among the nucleosomes can be distinguished by classification and visualized by showing different categories of nucleosomes (Figure 3A, indicated by different colors), suggesting that chromatin composed of different nucleosome categories may affect chromatin flexibility and can be observed by this strategy. Moreover, we noticed that if most of the nucleosomes fall into the same category (same color), the nucleosome oligos tend to adopt a typical tetranucleosome configuration. In contrast, the incorporation of an increasing number of poor nucleosomes in the nucleosome oligos would disrupt the regular packing and weaken the inter-nucleosomal interactions, leading to a progressively distorted and extended conformation and a reduced structural stability. Interestingly, we further noticed that the partial unwrapping of DNA in some nucleosomes would hinder the folding of the tetranucleosome and lead instead to a beads-on-a-string like structure (Figures 3B and S6B). These results indicated that the Nuc-back strategy could link nucleosome categories with chromatin architecture and help to analyze the chromatin at the single nucleosome level.

**Figure 3.**
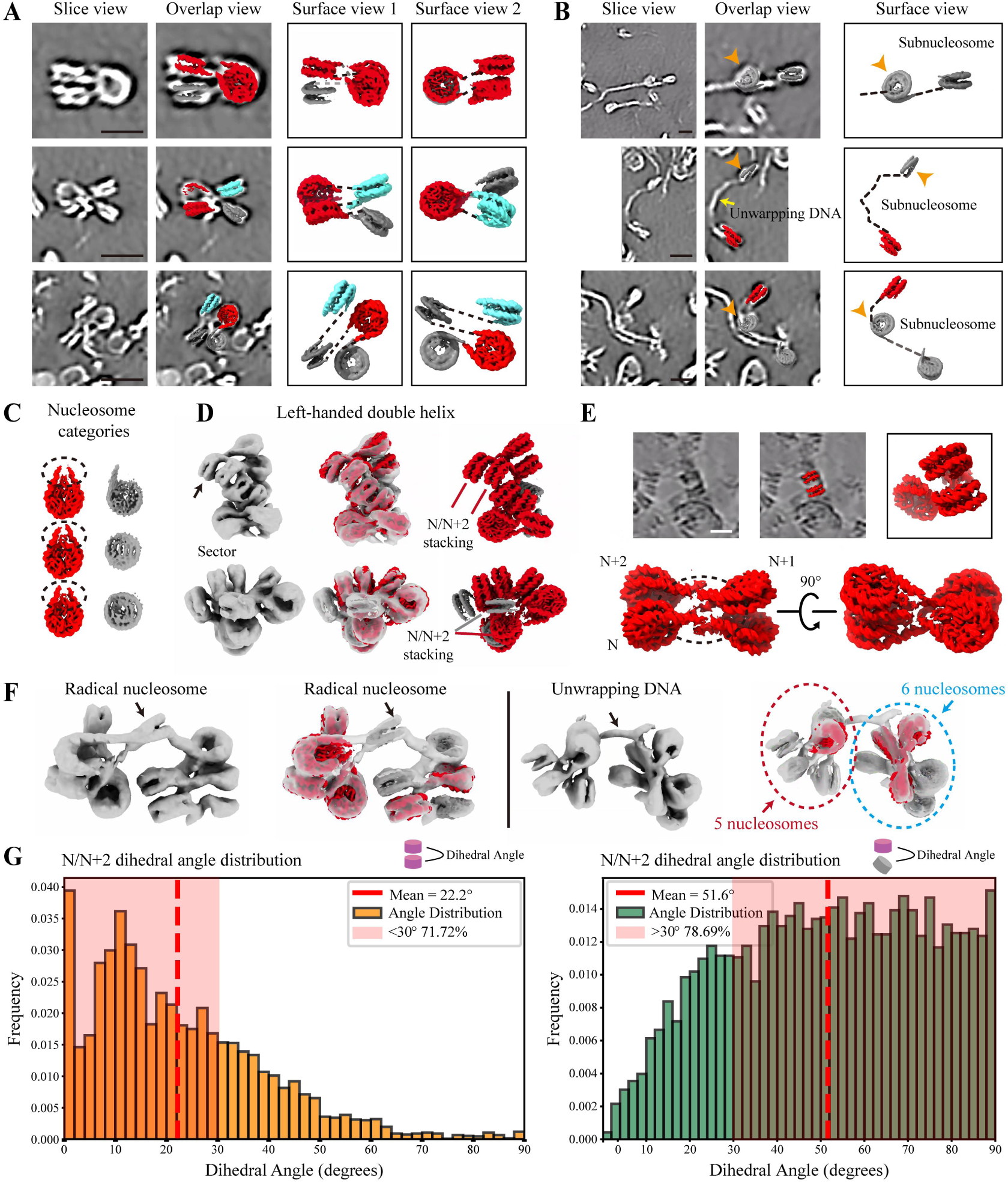
Morphology analysis of the 12× reconstituted chromatin fiber. **(A)** Visualization of tetranucleosomes. From left to right, each chromatin is presented with a tomographic slice, an overlay of the slice and Nuc-back map, Nuc-back map visualization, and a rotated view for Nuc-back map, respectively. Nucleosomes are connected by dashed lines based on the signal observed in the tomographic slices. **(B)** Visualization of sub-nucleosome averaged map and unwrapping of DNA in tomographic slice. Each region is presented with a tomographic slice, an overlay of the slice and Nuc-back map, and Nuc-back map visualization. Nucleosomes are connected by dashed lines based on the signal observed in the tomographic slices. **(C)** Classification categories of nucleosomes on the 12× reconstituted chromatin. The nucleosomes which the linker DNA densities resolved on both sides are colored red, radical nucleosomes are colored grey. **(D)** Morphology analysis of 12x chromatin fibers. Shown are the segmentation results, the overlay of segmentation with the Nuc-back map, and the Nuc-back map. The arrows indicate nucleosome density. **(E)** Connection between consecutive nucleosomes in chromatin revealed by Nuc-back analysis. Nucleosomes are connected to form a relatively typical tetranucleosome structure. **(F)** Two representative irregular chromatin fibers. The disruption of the regular DNA trajectory is caused by either radical nucleosome or DNA unwrapping in fibers. Segmentation and overlap views of two chromatin fibers are shown. **(G)** The distribution of dihedral angles between N/N+2 nucleosomes in constrained nucleosomes (left) and in constrained nucleosomes and radical nucleosomes (right) based on the Nuc-back analysis result. Scale bars:10 nm.

### Interpretation of local chromatin organization and interactions between nucleosomes

Beyond tetranucleosomes, we then applied the Nuc-back strategy to study longer chromatin fibers. For clarity, nucleosome categories in subsequent samples are displayed using three types: linker DNA densities resolved on both sides (red), linker DNA density resolved on only one side (blue), and a polymorphic category combining sub-nucleosomes and nucleosomes with high structural variations result in relatively abnormal density (gray).

In the reconstituted chromatin with 12× nucleosomes (dataset 1), to characterize the 3D structure, we obtained the 3D volume of the chromatin particle using DeepWedge [27], which simultaneously denoises the tomogram and compensates for the missing wedge. The 3D features of chromatin architecture revealed from the segmentation allow us to determine the number, orientation, and overall morphology of nucleosomes (Figure 3C, S8A). Some chromatin fibers exhibited left-handed helical and sector-shaped features, similar to those we observed in the previous single particle analysis [12], indicating that such structural motifs represent a fundamental and widespread mode of chromatin organization. Nevertheless, it was insufficient to resolve the conformational state of individual nucleosome, which motivated us to apply the Nuc-back strategy. The Nuc-back analysis showed that the tetra-nucleosome-like subunits and the nucleosome stacks between the N/N+2 nucleosomes could be observed frequently (Figure 3D, 3E). At single nucleosome level, most nucleosomes (red) showed linker DNA densities on both sides, suggesting a relatively confined relation between N/N+1 nucleosomes. We defined nucleosomes in this category as constrained nucleosomes (red) and the others as radical nucleosomes (gray). Interestingly, within the constrained category, the nearly-touching linker DNA densities allowed a reliable trace of the DNA trajectory between sequential nucleosomes (Figure 3E). The result showed that nucleosomes are arranged in an unambiguous zig-zag pattern, consistent with our previous structure [12]. This result indicated that the Nuc-back strategy can well trace the connection between nucleosomes based on the resolved linker DNA densities in some specific chromatin regions.

Interestingly, we also observed some irregularly chromatin fibers whose arrangement of nucleosomes is different from the others (Figures 3F and S8B). By observing different categories of nucleosome arrangements, we found that the radical nucleosome or DNA unwrapping can disrupt the regular trajectory of DNA along the chromatin fiber. In one striking case, we identified eleven nucleosomes in the 12× reconstituted chromatin, but a gap was created by a stretch of free DNA. Consequently, one nucleosome was missing and an irregular DNA path was formed (Figure 3F). The DNA unwrapping was accompanied by free DNA densities surrounding the nucleosome particle (Figures S8B and S8C). Even in the regular chromatin regions, while tracing DNA among nucleosomes, we found the DNA trajectories in the radical nucleosomes appear much less clear, indicating that the relative positions of these nucleosomes to their neighborhoods (i.e., N/N+1) were less constrained (Figure S8D). These observations suggest that the radical nucleosome can disrupt the DNA trajectory, representing an important source of chromatin irregularity.

Furthermore, different from the constrained category of nucleosomes which facilitated the nucleosome stacking, the radical nucleosomes also disturbed N/N+2 nucleosome stack formation, often leading to an irregular path at that site along with the chromatin extension (Figures 3C and S8A). The statistical analysis of the mean dihedral angles between N/N+2 nucleosomes also confirmed this observation. As shown in Figure 3F, a better stacking was showed around the constrained nucleosomes (mean dihedral angle: 22.2°, proportion <30°: 71.72%), whereas the stacking involving around radical nucleosome category was significantly worse (mean dihedral angle: 51.6°, proportion >30°: 78.69%). Taken together with the results from the tetranucleosomes, these findings indicate that nucleosome heterogeneity contributes to chromatin irregularity in multiple aspects.

Apart from the reconstituted chromatin, we also applied the same strategy to the native chromatins, including those extracted from HEK293F cells (dataset 5, dataset 6) and the *in vivo* chromatins within cryo-lamella of frog erythrocyte (dataset 3). Compared to the 12× reconstituted fibers, due to heterogeneous DNA and histone composition, the nucleosome classes are more diverse, and the native chromatin appears much more irregular (Figures 1D and S9A). In spite of that, a general zig-zag pattern can still be identified and frequently observed, particularly in the loosely packed regions (Figures 4A and 4B). These results indicated that, even with the increased flexibility, the nucleosomes in the native chromatin remain in a zig-zag arrangement, consistent with the reconstituted chromatin fibers. However, some nucleosomes are found unexpectedly far from their neighbors in the native chromatins, which may be resulted from a long nucleosome repeat lengths (NRL). This would further enhance the local flexibility of chromatin in which the nucleosomes become much more dynamic in the unconstrained environment (Figures 4B and 4C)

**Figure 4.**
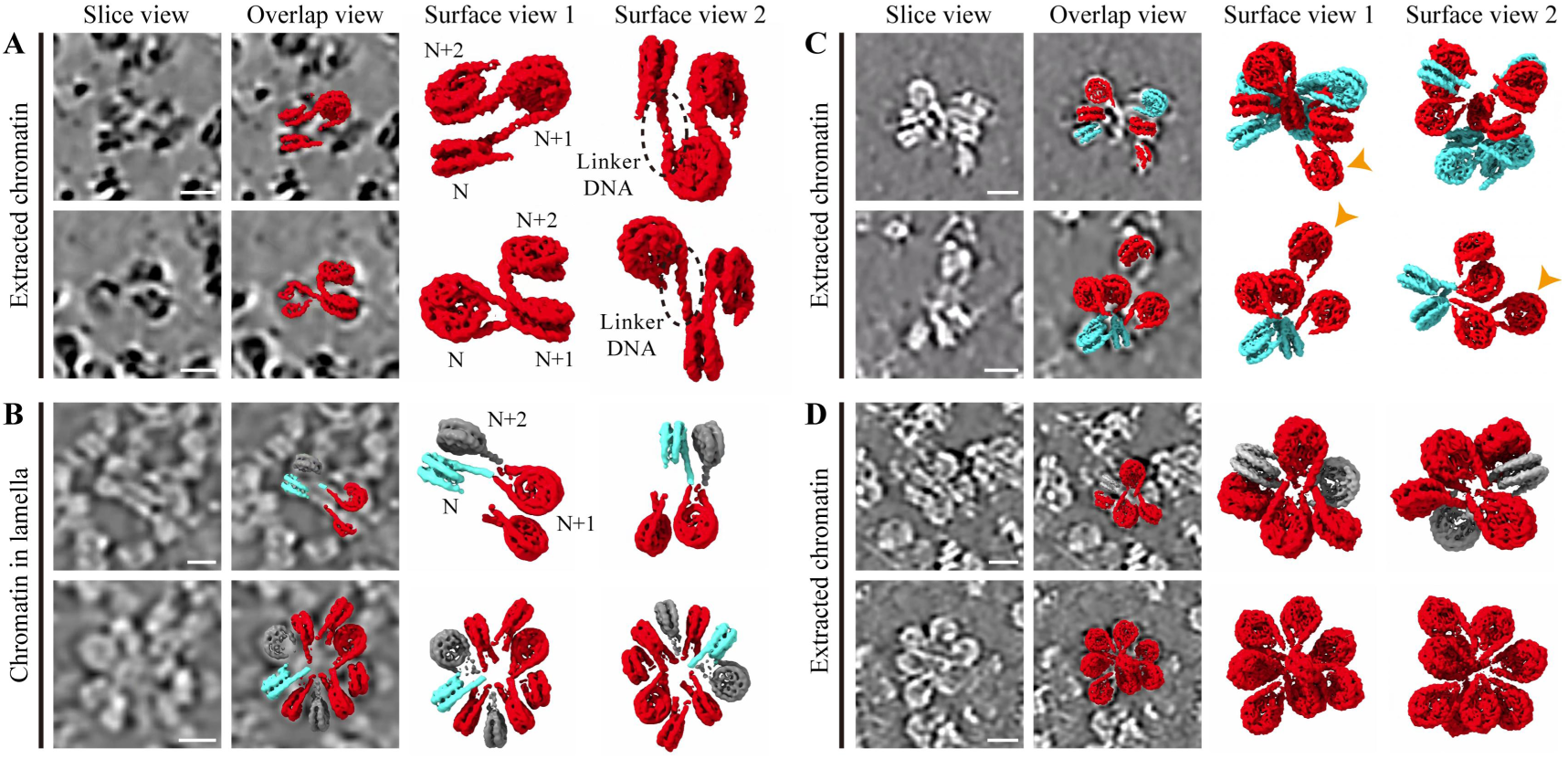
Polymorphism of native chromatins. **(A)** Chromatin particles extracted from HEK293F cell. Nucleosome are connected by the STA averaged density of the linker DNA, demonstrating a zig-zag DNA trajectory. **(B)** Visualization of *in vivo* chromatins within the cryo-FIB lamella of frog erythrocyte, demonstrating a typical zig-zag configuration. **(C)** Chromatin fibers extracted from HEK293F cell. The nucleosome marked with an orange arrow is far away from other compacted nucleosomes, likely due to a longer NRL. **(D)** Extracted chromatin fibers from HEK293F in high salt concentration show a highly compact configuration. For A-D, in each panel, the chromatin is presented with a tomographic slice, an overlay of the slice and Nuc-back map, Nuc-back map, and a rotated view for Nuc-back map, respectively.

Interestingly, although the heterogeneous compositions of native chromatin comprised diverse nucleosome structures, we fortuitously identified regions dominated by homo-categories under specific conditions (Figure 4D). For instance, when the extracted chromatins (dataset 6) were prepared under relatively high salt concentration, they displayed high compaction and the nucleosomes were found arranged in a petal pattern with interdigitated linker DNA at the center, consistent with the previously proposed cross-linker model [28]. Nucleosomes in this tightly packed chromatin were predominantly classified into the constrained nucleosome category with linker DNA on both sides and exhibited a high proportion of H1 binding—similar to the 12× reconstituted compact chromatin fibers (Figures 3D and S9B). In contrast, in some loosely packed fibers, H1 may fail to bind to the linker DNA region, resulting in a single-sided linker DNA class (Figures 3B and 3C). Hence, the binding ratio of H1 to nucleosomes also regulates the morphology and regularity of chromatin.

Above all, by visualizing the chromatin at individual nucleosome level, we are able to analyze the underlying factors of chromatin flexibility, e.g., heterogeneity of nucleosomes, different NRL, and the presence or absence of linker histone. These factors in turn would affect the nucleosome stacking and the spacing between nucleosomes, thereby contributing to the flexibility of chromatin.

### Visualization of the architecture between regulators and chromatin

Chromatin regulators, e.g., protein factors introduce histone or DNA modifications or chromatin remodeling, play crucial roles in regulating gene expression. In recent years, various structures of *in vitro* reconstituted nucleosome in complex with different chromatin regulators have been solved by cryo-EM single particle reconstruction. However, how chromatin regulators bind to poly-nucleosomes and particularly the *in vivo* chromatin remains unclear, likely due to the difficulty of averaging the flexibility across nucleosomes. In this study, we used the binding of histone acetyltransferase (HAT) to chromatin as a model to explore if the Nuc-Back strategy could be applied to reveal the binding of chromatin regulators to chromatin, e.g., to locate the HAT-bound nucleosomes in the context of chromatin (Sampling details in Materials and methods).

First, we mixed a purified HAT complex, piccolo NuA4 (pNua4), and chromatin extracted from HEK293F cell at a HAT: nucleosome ratio of 3:1 (dataset 7, Figure 5A). Through STA and classification, the nucleosomes were categorized into approximately six distinct classes, as indicated in Figure 5B. In two of them, extra density could be clearly observed on the disc surface of nucleosome, either on one side or both sides. The densities of these two classes can be well fitted with the structures of nucleosome with H1 (PDB:7XBP) and NuA4 (PDB:8X2X) (Figure 5C). NuA4 was found to bind to chromatin in a way similar to that with single nucleosome, in consistent with the previously solved structures of NCP-NuA4 complex of by SPA [5].

**Figure 5.**
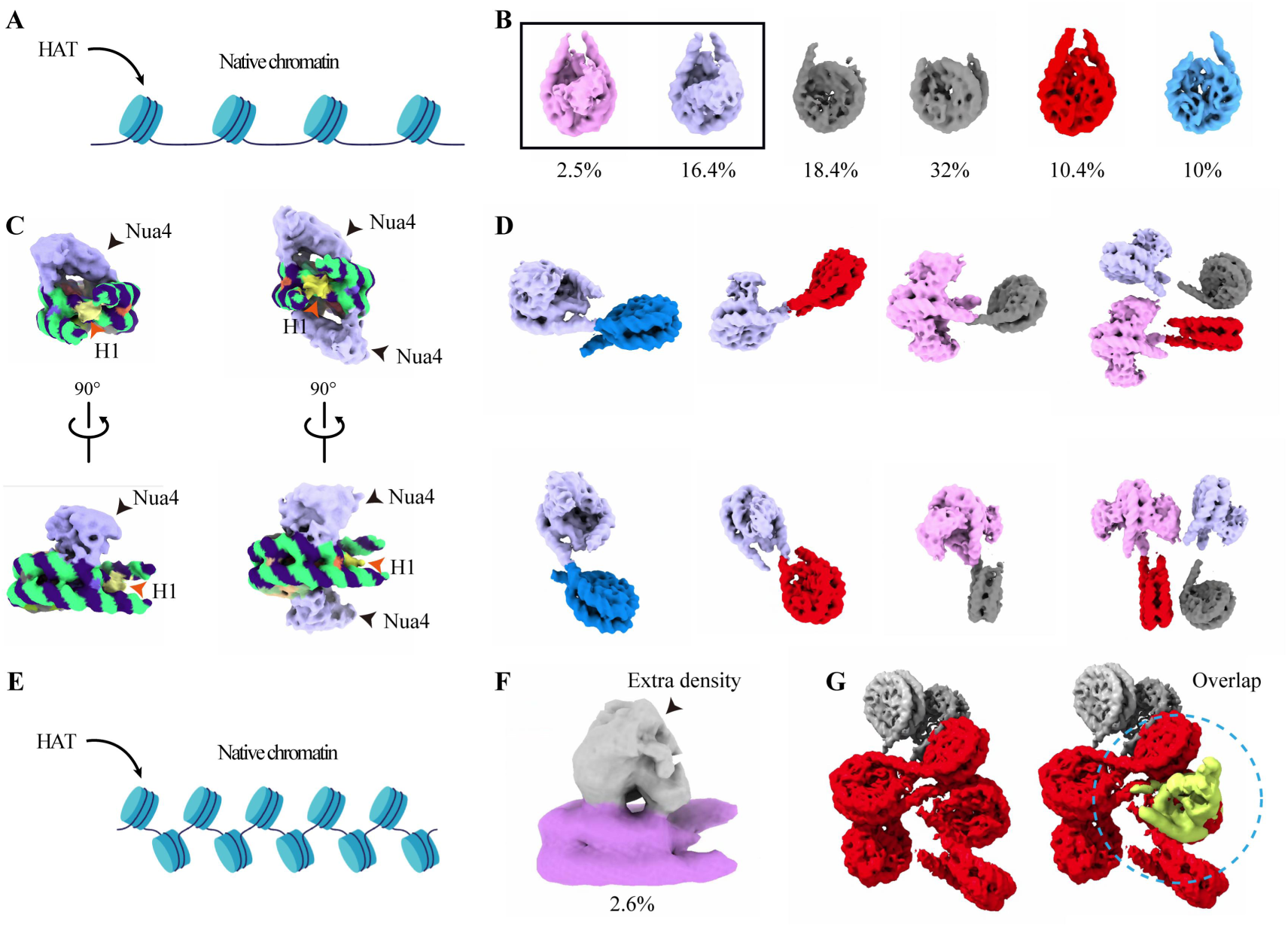
Locating the regulators in chromatin. **(A)** Schematic diagram of the sample preparation, adding HAT to the loose extracted chromatin. **(B)** Different nucleosome classes obtained by STA and classification. **(C)** Two of the six classes in (B) showing the nucleosomes in complex with Nua4 (one side and two side). The Nua4 density is shown in purple and marked with arrows. **(D)** Structure of the multi-nucleosome assembly with Nua4 proteins, reconstructed from different mapped-back nucleosome categories. **(E)** Schematic diagram of the sample preparation, adding HAT to the compact extracted chromatin. **(F)** Structure of the HAT-bound nucleosome from compact extracted chromatin, with the extra density of HAT marked with arrow. **(G)** Visualization of chromatin reconstructed by Nuc-back without (left) and with (right) the density of HAT-bound nucleosome class (highlighted in green) mapped back into the chromatin fiber density.

Next, we used the Nuc-back strategy to display the HAT-bound nucleosome categories within the chromatin map. We found that the chromatin became relatively loosen upon the binding of NuA4 compared with the original folding state (Figure 5D), possibly caused by the NuA4 binding across multiple consecutive nucleosomes. Binding to the disc surface of nucleosome, NuA4 interfered sterically the interaction between N/N+2 nucleosomes, preventing local nucleosome stack formation (Figure 5D). Interestingly, we observed that NuA4 can bind a single nucleosome on chromatin in a 1:2 manner, with both disc surfaces of the nucleosome capable of interacting with NuA4 (Figure 5D). These results suggest that the Nuc-back strategy can visualize the interplays between poly-nucleosomes and chromatin regulators, e.g., NuA4, enabling a better understanding of how chromatin regulators bind across multiple nucleosomes. Meanwhile, we can also observe that regulatory factors occupy certain spatial regions, thereby influencing chromatin structure.

To assess if the Nuc-back strategy could locate the chromatin-bound proteins with lower-abundance, we mixed HAT complex and chromatin at a lower HAT: nucleosome ratio of 0.5:1 (dataset 8). Given the low binding ratio, it is challenging to distinguish HAT-bound nucleosomes from the large number of unbound particles directly via 3D classification. To do this, we first defined the fundamental nucleosome categories based on the STA of nucleosomes at a moderate-to-high resolution and then applied Nuc-back strategy to ensure accurate nucleosome orientation assignment (Figure S9C). The resulting chromatin map appeared more compact than above. We then performed iterative classification focused on the nucleosome periphery to identify sub-category which shows nucleosome with extra HAT complex density (Figure 5F). Mapping the HAT-bound nucleosome complex back into the chromatin map, we found that, despite the lower binding frequency (2.6%), the HAT density spatially overlapped with a specific nucleosome can be well identified in the compact chromatin (Figure 5G). This result demonstrates that the overlay-based strategy could be able to accurately locate the protein-bound nucleosomes in the chromatin context, even in a low abundance.

## Discussion

Chromatin is a dynamic, high order assembly of nucleosomes, enabling the packaging of genomic DNA into the confined space of the nucleus. However, structural analysis of such irregular assemblies remains challenging due to their intrinsic flexibility. The conformational diversity of chromatin poses a significant challenge for conventional averaging-based reconstruction methods, which appear inadequate for resolving such flexible assemblies. Moreover, the underlying correlation between single nucleosome heterogeneity and chromatin polymorphism has remained poorly understood. In this study, we achieved medium-to-high resolution (9.5-4.1 Å) STA structures of nucleosomes from diverse samples. This improvement in resolution is crucial for subsequent nucleosome classification and mapping-back, as it enables fine structural categorization and accurate alignment, thereby ensuring precise determination of nucleosome center coordinates and orientations within the 3D chromatin map. Based on this advance, we established a Nuc-back strategy, which regards nucleosome categories as versatile building motifs to construct irregular chromatin architecture.

In contrast to prior studies, we present an extensive classification of nucleosomes. Our study showed that increasing the number of classes had two advantages: first, it reduced potential errors in chromatin reconstruction caused by insufficient resolution or mis-oriented nucleosomes; second, it allowed the separation of distinct nucleosome categories with different structural features, such as variations in the histone region or linker DNA geometry, the presence or absence of linker histone or chromatin regulators. By mapping back each individual nucleosome with the corresponding nucleosome category density, we found that almost all of the nucleosome densities in chromatin map could be well assigned. Notably, the nearly-touching linker DNA densities between two adjacent nucleosomes allowed to trace the trajectory of these nucleosomes in chromatin. This makes Nuc-back a powerful tool that not only provides the overall chromatin architecture (the spatial arrangement of nucleosomes) but also displays the structural state of each individual nucleosome within chromatin.

To reveal the correlation between single nucleosome heterogeneity and chromatin polymorph, we used Nuc-back to analyze the interaction among nucleosomes within the chromatin. First, we used the reconstituted chromatin to study its intrinsic flexibility. In general, we found that the structure of individual nucleosome component is heterogeneous, and the states of nucleosomes within the chromatin are also variable. This suggests that nucleosome composition heterogeneity may be a major contributing factor to the overall heterogeneity of chromatin. Our study of the 12× reconstituted chromatin revealed that for those nucleosomes falling in the well-constrained categories, we were able to reliably trace the zig-zag trajectory of the linker DNA, and the stacking between N and N+2 nucleosomes is well maintained. However, if one or more radical nucleosomes present in the chromatin, they may induce global or local irregularity in the DNA trajectory. As a result, the trajectory of the linker DNA could not be precisely followed, and the stacking angle between N and N+2 became larger, producing a heterogeneous chromatin architecture. Thus, our results indicate that although the two-start zig-zag appears as a generally local folding pattern, the variability in nucleosome states introduces significant heterogeneity into the overall chromatin fiber.

Our analysis of extracted and *in situ* nucleosomes showed that they are significantly more heterogeneous than those in the reconstituted chromatin. Early studies on the nucleosome dynamic indicated that linker histone binding and special DNA sequence (i.e. 601 DNA) contributed significantly to nucleosome stabilization [29]. In line with that, our results showed the uneven distribution of linker histone and native DNA sequence in the extracted and *in situ* nucleosomes could make the nucleosomes less stable and increase the overall flexibility of the chromatin fiber. Studies on the nucleosome NRL suggested that NRL variation had significant impacts on the nucleosome stacking and chromatin folding [16,30,31]. Consistently, in the present study, we found that longer NRLs led to sudden increases in the distance between adjacent nucleosomes, which unfaverating nucleosome stacking thus rendering local polymorphic chromatin architecture.

The chromatin regulators play an important role in the regulation of gene expression. However, investigating their interactions on multi-nucleosomes remains challenging due to structural heterogeneity. Taking the histone acetyltransferase NuA4 as an example, we studied chromatin regulators binding to chromatin using the Nuc-back strategy. Our results demonstrate that Nua4 localizes within native multi-nucleosomes in a manner dependent on nucleosome disc exposure. We found that binding of NuA4 to a nucleosome surface prevents other nucleosomes from approaching that surface. When NuA4 bound between N/N+2 nucleosomes, steric hindrance increased their separation, creating more open chromatin space in that region. Additionally, we demonstrated that with low abundance binding proteins, Nuc-back strategy is still feasible to classify and localize protein-bound nucleosomes in compact chromatin. This approach provides insight into the spatial organization of protein binding in multi-nucleosome contexts and helps elucidate how chromatin architecture influences protein localization. Based on the observation with the Nuc-back strategy, the sources of chromatin flexibility could be spanned from nucleosome heterogeneity, to nucleosome interactions, and finally to flexible chromatin.

In summary, by improving the STA averaged resolution of nucleosome in the context of chromatin, we established a Nuc-back strategy utilizing classified nucleosome categories as building motifs to reconstruct irregular chromatin architecture. The Nuc-back strategy enables single-nucleosome-level chromatin visualization, tracing trajectories and distinguishing heterogeneity in both histone and DNA regions. Applying this strategy across chromatin revealed that local nucleosome heterogeneity primarily affects chromatin structure by regulating the DNA trajectory and N/N+2 interactions, elucidating the correlation between nucleosome heterogeneity and chromatin polymorphism. Moreover, Nuc-back could be used to visualize the specific binding mode of chromatin regulators to chromatin, even at low abundance. Our findings illuminate general chromatin architecture and specific nucleosome interactions, providing a powerful tool for future studies on 3D genome organization and epigenetic regulating process.

## Materials and methods

### Reconstitutions of histone octamers, nucleosomes and chromatin fibers

Recombinant wild-type *Xenopus laevis* histones (H2A, H2B, H3, and H4) were expressed in *E. coli* BL21(DE3) cells and purified as previously described [12]. DNA fragments containing tandem repeats of either 4×177 bp or 12×177 bp 601 sequence for chromatin reconstitution were cloned and purified as described previously [12].

Histone octamers were reconstituted as previously described. Briefly, equimolar amounts of H2A, H2B, H3 and H4 were mixed in unfolding buffer (7 M guanidinium HCl, 20 mmol·L^-1^ Tris-HCl, pH7.5, 5 mmol·L^-1^ 2-mercaptoethanol), and then dialyzed against refolding buffer (2 mol·L^-1^ NaCl, 10 mmol·L^-1^ Tris-HCl, pH 7.5, 1 mmol·L^-1^ EDTA, 5 mmol·L^-1^ 2-mercaptoethanol). The resulting histone octamers were purified through HiLoad 16/600 Superdex 200 pg column (GE Healthcare, USA), and peak fractions were collected and stored.

Nucleosome arrays and 30-nm chromatin fibers were assembled using the salt dialysis method as previously described [12]. Histone octamers and 4×177 bp or 12×177 bp 601 DNA were mixed at equimolar ratios in refolding buffer, and was continuously diluted by slowly adding TE buffer (10 mmol·L^-1^ Tris-HCl, pH 8.0, 1 mmol·L^-1^ EDTA) over 16 h at 4°C to reduce the NaCl concentration from 2 mol·L^-1^ to 0.6 mol·L^-1^. Then, H1.4 was added at an equimolar ratio relative to mono-nucleosomes, and the mixture was further dialyzed against TE buffer containing 0.6 mol·L^-1^ NaCl for 3 h, followed by a final dialysis against HE buffers (10 mmol·L^-1^ HEPES, pH 8.0, 0.1 mmol·L^-1^ EDTA) for 4 h.

### Preparation of the native chromatin

HEK293F cells were maintained in suspension culture with 5% CO_2_ at 37℃ in FreeStyle^TM^ 293 expression medium (Thermo Fisher Scientific, USA) and supplemented with 1% (v/v) fetal bovine serum (FBS). Sf9 insect cells were maintained in suspension culture at 27℃ in serum-free sf-900^TM^ Ⅱ SFM medium (Thermo Fisher Scientific, USA). For chromatin extraction, HEK293F and Sf9 cells were expanded to a density of 1.8-2.0 × 10^6^ cells/mL in a final volume of 200 mL. Cells were harvested by centrifugation, washed twice with phosphate buffered saline (PBS), and resuspended in 20 mL Lysis buffer (0.25 mol·L^-1^ sucrose, 60 mmol·L^-1^ KCl, 15 mmol·L^-1^ NaCl, 10 mmol·L^-1^ MES, 5 mmol·L^-1^ MgCl_2_, 1 mmol·L^-1^ CaCl_2_, 0.5% Triton X-100, 0.1 mmol·L^-1^ PMSF, pH6.5) at 4℃. The cell suspension was homogenized by 30 strokes of a Dounce homogenizer over 10 min on ice. Nuclei were pelleted by centrifugation at 4,000 rpm for 5 min at 4℃, and resuspended in 2 mL digestion buffer (10 mmol·L^-1^ PIPES, 2 mmol·L^-1^ CaCl_2_, 10 mmol·L^-1^ NaCl, pH 8.0).

The isolated nuclei were warmed at 37℃ for 5 min, and MNase (1:1000 dilution) was added to the suspension and incubated for 15 min at 37℃. Digestion was stopped by adding 10 mmol·L^-1^ EDTA, followed by rapid cooling on ice and centrifugation at 10,000 rpm for 5 min at 4℃. The supernatant containing soluble native chromatin was collected, and DNA concentration was determined by measuring absorbance at 280 nm with a NanoDrop 2000C spectrophotometer (Thermo Fisher Scientific, USA).

Chromatin samples were carefully loaded onto a 5 mL 5-40% sucrose gradient prepared in TE buffer and centrifuged in a Beckman MLS-50 rotor at 35,000 rpm for 3 h at 4℃. Fractions enriched in particles containing DNA fragments corresponding to 6-12 nucleosomes were collected, divided into two equal portions, and dialyzed for 48 h at a sample-to-buffer ratio of 1:500 against either HE buffer 1 (10 mmol·L^-1^ HEPES,10 mmol·L^-1^ NaCl, 0.1 mmol·L^-1^ EDTA, pH=7.5) or high-salt HE buffer 2 (10 mmol·L^-1^ HEPES,100 mmol·L^-1^ NaCl, 0.1 mmol·L^-1^ EDTA, pH=7.5). Nucleosomal DNA fragment sizes were determined by agarose gel electrophoresis after deproteinization with 1% SDS and 0.5 mg/mL proteinase K.

### Sample Preparation and vitrification for cryo-ET

The HAT complexes were expressed in *E. coli* following the previous protocols [5], mixed with native chromatin under two buffer conditions (10 mmol·L^-1^ NaCl and 100 mmol·L^-1^ NaCl) at molar ratios of 3:1 and 0.5:1 respectively., and followed by incubation at 4℃ in 10 mmol·L^-1^ HEPES, 50 mmol·L^-1^ NaCl, pH 7.0 for 30 min. The HAT-chromatin complex was loaded on a 5 mL 5-40% sucrose gradient prepared in HE buffer 3 (10 mmol·L^-1^ HEPES, 10 mmol·L^-1^ NaCl, 0.1 mmol·L^-1^ EDTA, pH=7.0) and centrifuged in a preparative ultracentrifuge with a Beckman MLS-50 rotor at 35,000 rpm for 3 hrs at 4℃. Fractions were collected and checked by 15% SDS-PAGE and agarose gel electrophoresis. Guided by the result, the fractions containing the homogeneous complex were combined, the glycerol was removed, and the complex was concentrated to 100 ng/μL.

Vitrobot Mark IV (FEI, USA) was used to prepare the reconstituted chromatin and extracted chromatin. 3.5 µL samples were applied to the plasma-cleaned Quantifoil R2/2 holey Cu grids with a blot force of 3 for 3 s in 100% humidity. Grids were then plunged into liquid ethane, which was cooled by liquid nitrogen.

### Cryo-lamella preparation using cryo-FIB

For the *in vivo* study, the frog erythrocyte nuclei were chemically fixed by 0.5% glutaraldehyde and 1% paraformaldehyde. The nuclei were further cryo-protected by glycerol at a final concentration of 3%. Aliquots of 1.5 μL sample (∼ 700 cells) were applied onto glow-discharged Quantifoil R2/1 300 mesh holey carbon grids, incubated for 10 s at 37°C and 20% humidity, blotted for 8 s with a filter paper and then plunged into liquid ethane using an EM GP systern (Leica Microsystems, Germany).

Vitrified cells were further processed by cryo-FIB milling for the preparation of lamellae. A dual-beam microscope FIB/SEM Aquilos 2 (Thermo Fisher Scientific, USA) equipped with a cryo-transfer system (Thermo Fisher Scientific, USA) and rotatable cryo-stage cooled at −191℃ by an open nitrogen circuit was used to carry out the thinning. During the cryo-FIB milling process, the milling angle between the FIB and the specimen surface was set to 5-10°. The milling was performed parallel from two sides to produce vitrified cell lamella. The accelerating voltage of the ion beam was kept at 30 kV, and the ion currents were in the range from 0.43 nA to 40 pA. The rough milling utilized a strong ion beam current of 0.43 nA and the final fine milling was operated with a small ion beam current of 40 pA. The thickness of the residual thin lamella with a good quality was < 150 nm.

### Cryo-ET data acquisition

For reconstituted chromatin and extracted chromatin (dataset 1, dataset 2, dataset 4-8), tilt series were collected on a Titan Krios cryo-TEM (Thermo Fisher Scientific, USA) at 300 kV with a BioQuantum energy filter operating at zero loss with a 20-eV slit width, and a K2 summit direct director (Gatan). The energy filter was used to increase image contrast. All tilt series used in this study were collected using a dose-symmetry strategy-based beam-image-shift facilitated acquisition scheme, by in-house developed scripts within SerialEM software. Low-magnification were taken at 8700x (object pixel size 1.69 nm) to obtain hole overviews. High-magnification tilt series were collected, operating in low dose mode. Tilt series were taken with a 3° tilt increment with an angular range from ∼−50° to 50°, with dose-symmetric tilt scheme. The K2 camera operating in dose fraction counting mode, recorded frames every 1 s for 3 electrons/Å² per tilt. The cumulative dose was in the range of 99 electrons/Å² per tilt series. For chromatin in lamella (dataset 3), 199 tilt series were collected on the same equipment. Tilt series were taken with a 3° tilt increment with an angular range from −60° to 40° starting at −10° to compensate for the lamella pre-tilt, with dose-symmetric tilt scheme. Targeted defocus, pixel size and other detailed parameters are listed in Supplemental Table 1.

### Tomogram reconstruction

All fractioned movies were performed using the Warp pipeline [32], including motion correction, contrast transfer function (CTF) estimation and creating stack. The tilt series were then automatically aligned using AreTOMO 1.1.0 [33], while the DarkTol parameter was adjusted to disable automatic removal of dark or low-quality tilts. Low-quality tilt series (e.g., dark or large shifts) were manually identified and excluded based on visual inspection on aligned tilt series, and their file numbers were recorded for reference. The iterative process continued with AreTOMO alignment until no further tilt series removal is needed. The alignment parameters for the remaining tilt series were transferred back to Warp, where initial tomogram reconstruction was performed at a pixel size of 10.88 Å for all chromatin datasets. Each dataset was denoised using three approaches: (1) training on a per-dataset and denoising using CryoCARE 0.3 [34], (2) training on the entire dataset and denoising using Warp Noise2Map [32], (3) training on a per-dataset and denoising and missing wedge reconstruction using DeepDeWedge 1.0 [27]. The method yielding better denoising performance was selected for final data presentation.

### Particle picking

For nucleosome picking within chromatin, we generated a 10.88 Å density map of a 147 bp nucleosome, which was subsequently low-pass filtered to 60 Å and used as the reference for template matching in the same pixel size tomograms. We initially performed particle picking using Gapstop [35], where nucleosomes were sampled every 5° to generate an angle list for template matching. A wedge list was generated based on the XML metadata files from Warp. Template matching was then carried out, and particles with a score above 0.08 (scores_threshold) were selected. These particles were subsequently cleaned using clean_by_otsu, yielding an approximate count of nucleosomes in the dataset. The selected coordinates and initial orientations were exported as a .star file.

For STA, we compared the initial orientations derived from template matching using pyTom [36], Gapstop (TM) [35], and Warp 1.0.9 [32]. Among these, Warp-based template matching provided more accurate initial orientations and identified a greater number of nucleosomes. Therefore, we performed final template matching in Warp. Based on the estimated number of nucleosomes in different datasets from Gapstop, we increased this number by 500–1000 and used it as the input for the “Store best results” function in Warp. Particle diameter was set to 110 Å and angular sampling was set to 15° (angle subdivisions). The resulting particle coordinates and orientations were then imported into RELION for further processing.

### Subtomogram averaging

For all chromatin datasets, particles were first extracted in Warp with a pixel size of 5.44 Å and a box size of 40. Initial cleaning was performed using 3D classification using initial orientations from Warp with local angle search in RELION 4.0 [25] to remove junks. To mitigate potential model bias, all orientation information was then discarded, and a global refinement from scratch was performed, which further eliminated low-quality particles (Fig. S1). For chromatin in lamella, two additional steps were included in the data processing of this dataset. Firstly, to reduce false positives, we manually defined the boundaries of the lamella in Dynamo 1.1.558 [37] and excluded regions above and below the lamella (Fig. S5). Secondly, the 3D particles were projected into 2D, followed by 2D classification to further remove low-quality particles.

The particles were then upsampled to 2.72 Å/pixel with a box size of 80. For these all-nucleosome particles, an elliptical mask was applied to conduct a global 3D classification (T=2, iteration=100) in RELION, which enabled multi-class visualization of Nuc-back. As for the reconstituted chromatin and extracted chromatin, after discarding poorly resolved classes, the dataset was upsampled to the original pixel size (bin1), and the nucleosome core was refined using a focused mask. As shown in Fig. S1, refinement was performed iteratively in both RELION and M [24]. In reconstituted chromatin, following several rounds of refinement, a final set of 38,321 particles was selected, yielding a nucleosome core structure with a reported resolution of 4.1 Å. With accurate orientations, we then performed a non-aligned 3D classification with a spherical mask focused on the H1 region, adjusting the T-value and mask diameter to resolve different H1 states. The resulting subclasses were further refined. In extracted chromatin, 138,850 particles were selected, yielding a nucleosome core structure with a reported resolution of 5 Å. Using a classification approach similar to the above with 91,312 selected particles, we obtained the structure of the nucleosome core particle with H1. For the nucleosome–HAT data, regardless of high or low binding ratios, we ignored the density of the associated proteins and focused on identifying the nucleosome itself. Based on the well-resolved nucleosome structure, we then performed classification specifically on the nucleosomal surface to identify classes that exhibited additional protein density. These classes were subsequently mapped back into 3D space for further structural analysis.

For chromatin in lamella, as shown in Fig. S5, after removing junk particles, 1,175,626 particles remained. We performed 3D classification on these particles to generate different classes; however, only 11,330 particles belonged to the class corresponding to the canonical nucleosome. We then performed C2 symmetry, resulting in a nucleosome reconstruction at 9.5 Å resolution. Subsequently, these particles underwent C1 refinement, followed by non-aligned 3D classification to resolve nucleosome classes with different H1 conformations. The selected subclasses were further refined to obtain the final structures.

The density maps were post-processed using either EMReady 1.3 [38] or the N2N denoising implemented in M [24].

### Nuc-back strategy optimization

We first optimized the Nuc-back strategy using cryo-ET data of *in vitro* assembled tetranucleosomes (dataset 4) (Figures 2A and 2B). By classification, we defined three categories of nucleosome (Figure 2D), including two categories in which the linker DNA densities were resolved on both sides (red) and on only one side (blue) respectively, and one category of poor resolved density (gray). After mapping these building motifs back to the tomographic slice and 3D segmentation (the denoising and segmentation were performed by using cryoCARE [34], Noise2Map [32] or DeepDeWedge [27]) with accurate center coordinates and orientation parameters, we found they aligned very well with the nucleosome densities in the tetranucleosome particles (Figure 2E). However, in spite of that most nucleosomes were fitted and assigned to one of the categories, some nucleosome-like densities remained unassigned to any category. We designated these as missing nucleosomes (Figure 2E).

To locate these missing nucleosomes, we examined the corresponding locations from the template matching step and explored the matching accuracy by visual evaluation. Briefly, if the center coordinates and orientation matched the slice feature, the parameters were recorded for further processing (Figure 2G). If the template-matching result were obviously discrepancy, manual refinement was performed and new parameters were saved (Figure 2H). A portion of nucleosomes (excluding junk particles) from previous classification were then combined with these newly identified nucleosomes, forming a new dataset (Figure 2I). This process facilitated assigning the missing nucleosomes to new class in another round of classification. After assigning all missing nucleosomes, the orientation of the averaged nucleosome density before and after refinement were compared (Figure 2J). If they are close enough, e.g., the angle variation was < 22.2°, the new averaged result was considered well refined and the newly averaged nucleosome density was mapped back using the new orientation (Figure 2K). If the angle variation > 22.2°, we consider the refinement was inaccurate for that nucleosome. 22.2° is derived from the mean dihedral angle distribution between nucleosomes (N/N+2) in 12x chromatin fibers. In such cases, we used either the former manually refined orientation or the initial template-matching result, ensuring all missing nucleosomes were properly located (Figure 2K).

### Measurement of the N/N+2 Dihedral Angle

After the Nuc-back analysis, different classes of nucleosomes were obtained. We defined two datasets based on classification: a set of well-structured nucleosomes and a set of poor-structured nucleosomes, each corresponding to a separate collection of .star files. Particles were further distinguished based on their tomogram IDs.

Using a home-made script, we computed the normal vector to the nucleosome plane by extracting orientation information from the .star files and applying it to the associated rotation matrices. For each well-structured nucleosome within a dataset, we searched for the nearest neighboring well-structured nucleosome within 10 nm and calculated the dihedral angle between their rotated normal vectors. If no neighboring nucleosome was found within 10 nm, the angle was not computed. This angle was defined as the stacking angle between well-structured nucleosomes.

Additionally, we introduced the poor-structured nucleosomes into the analysis. For each well-structured nucleosome, we searched for the nearest poor-structured nucleosome within 10 nm, and computed the dihedral angle between their respective normal vectors after rotation. This was defined as the stacking angle between well-structured and poor-structured nucleosomes. All angle measurements were performed in batch using custom scripts. The resulting dihedral angles were compiled across datasets and visualized as probability histograms to analyze and compare the stacking behaviors of nucleosomes across different classes.

### Visualization

The individual classes were refined in RELION using gold-standard refinement with independent half-set splitting. The resulting coordinates were then mapped back into the original tomograms using ArtiaX 0.1 [36], enabling visualization of the spatial distribution of each nucleosome within chromatin. To assess whether the nucleosome orientations and positions were consistent with the structural features observed in the tomographic slices, the denoised tomogram was overlaid with the mapped particles for visual inspection in ChimeraX 1.5 [39]. This allowed for a clear evaluation of the alignment between each nucleosome and the surrounding chromatin context.

## Supplementary data

Document S1. Figures S1–S10, Tables S1

## Acknowledgments

We would like to thank Xujing Li, Lulu Qin and Xiaojun Huang for their technical help and support with electron microscopy and cryo-FIB. All EM data were collected and processed at the Center for Bio-imaging (CBI), Institute of Biophysics (IBP), CAS.

## Author contributions

Conceptualization: P.Z., H.Z.; Formal analysis: H.Z., Y.L., C.P., F.B., C.Y., W.N., H.Y., K.S., P.Z.; Writing – original draft: P.Z., H.Z., Y.L.; Writing – review and editing: P.Z., H.Z., Y.L., K.S.; Visualization: H.Z., Y.L., C.P., H.Y.; Funding acquisition: P.Z., Y.L., K.S.

## Data availability

The subtomogram averaging results have been deposited in the Electron Microscopy Data Bank (EMDB): nucleosome core and H1-bound nucleosome from 12× reconstituted chromatin (EMD-66336, EMD-66335), nucleosome core and H1-bound nucleosome from SF9 cell chromatin (EMD-66337, EMD-66338), nucleosome core and H1-bound nucleosome from frog nuclei lamella (EMD-66340, EMD-66341).

