## Supplementary material for "Visualization of the architecture of flexible chromatin and the binding of chromatin regulators": Document S1

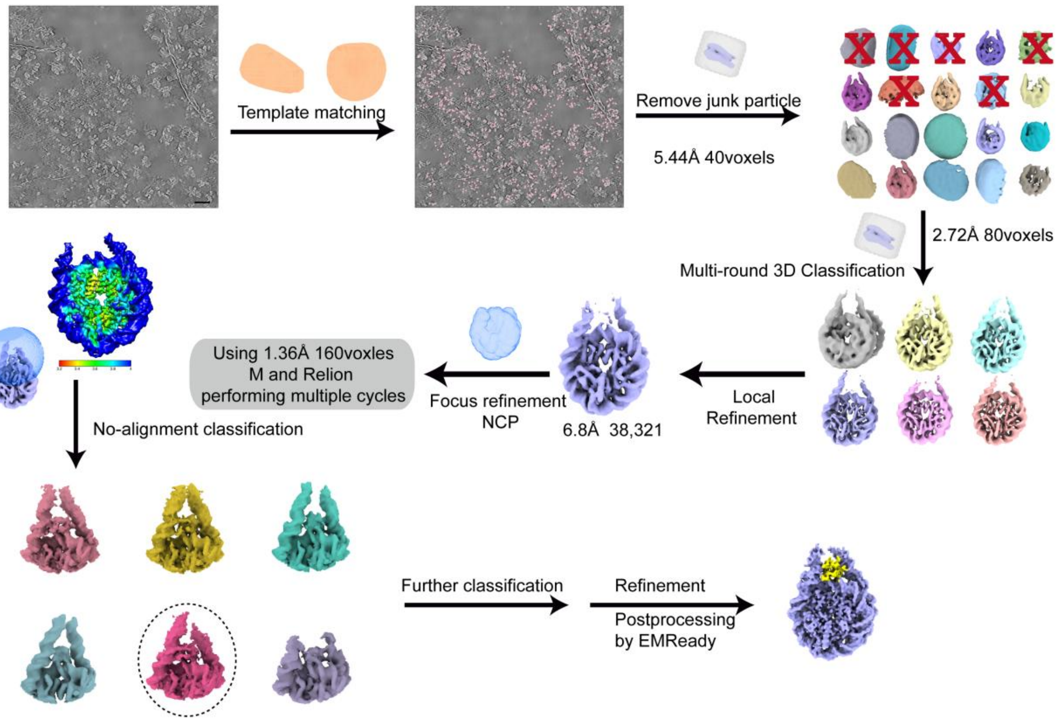


#### Figure S1. Flowchart of STA of nucleosomes in 12× reconstituted chromatin.

Nucleosomes in the reconstituted chromatin are initially selected by template matching. The selected nucleosomes are classified to remove junk particles, and subjected to further classification to improve the overall nucleosome resolution. After optimizing the nucleosome core, additional classification is performed for the H1 region to enhance its resolution. Scale bars: 30 nm.

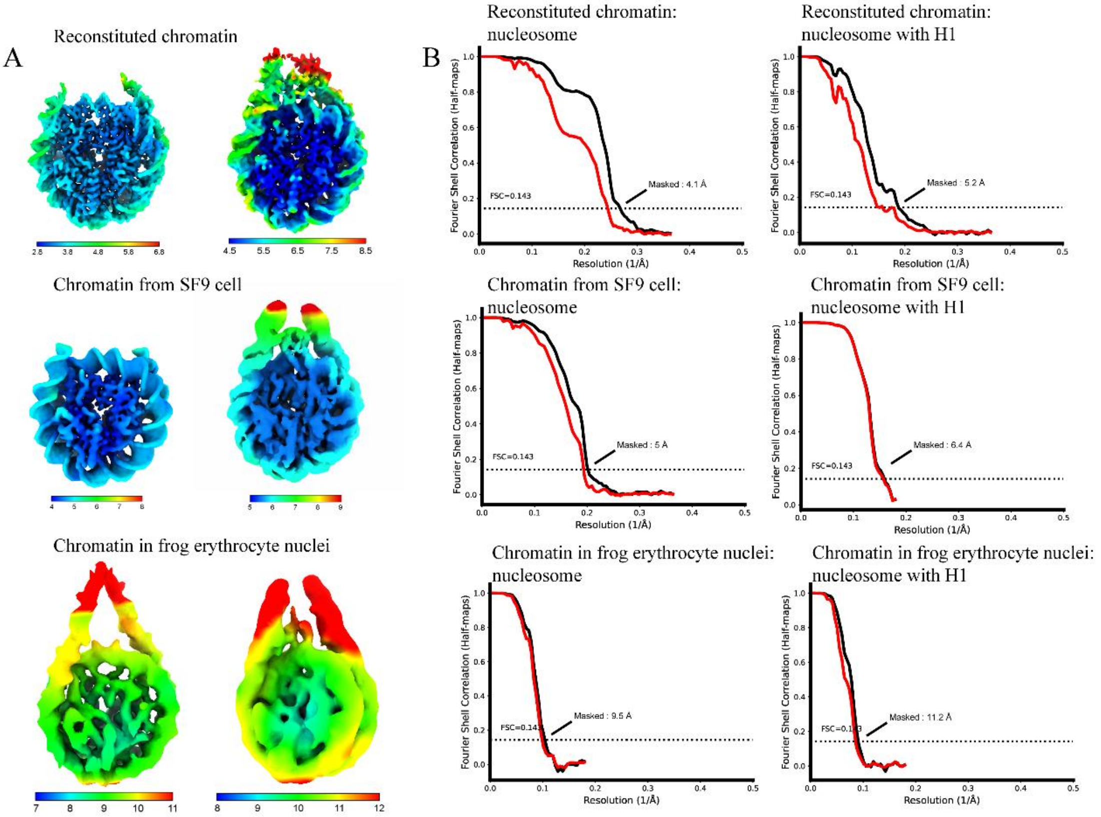


**Figure S2. Resolution characterization of 3D reconstruction of the nucleosome core particle and nucleosome with H1 from different chromatin source.**

(A) Local resolution map. (B) FSC curve.

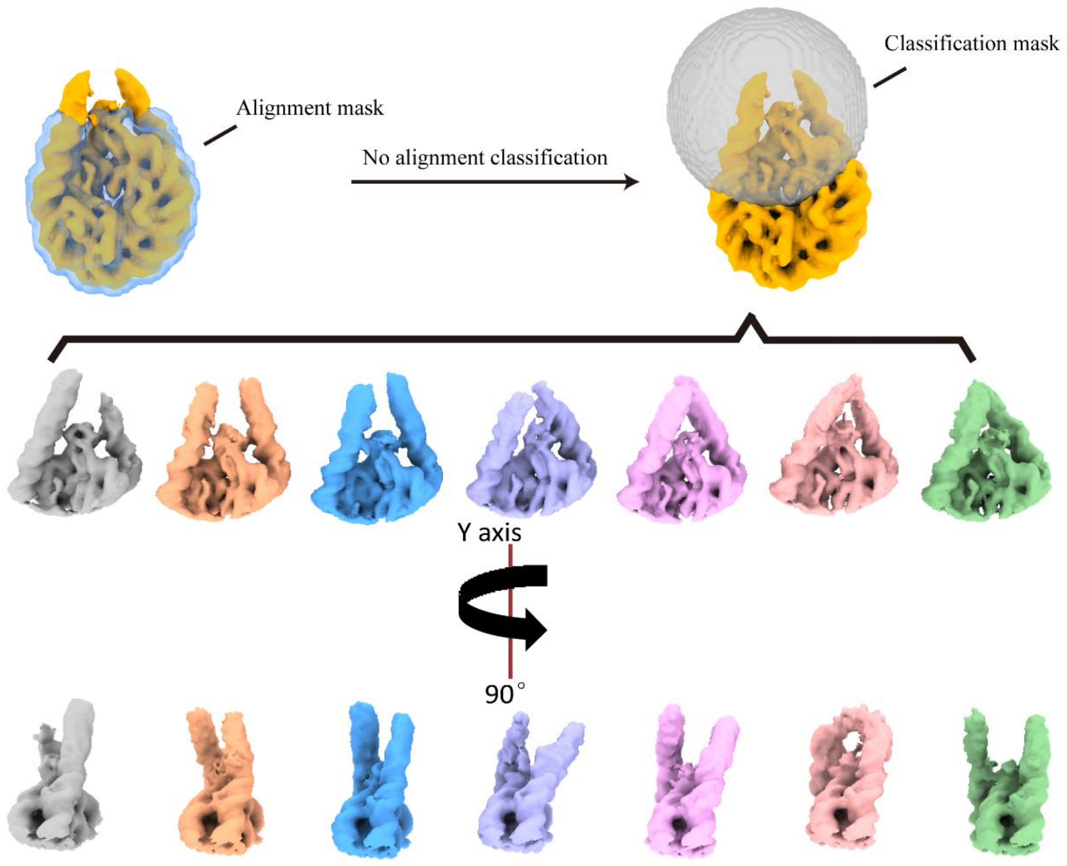


**Figure S3. Analysis of the dynamics for the chromatin fragments extracted from SF9 cells in nucleosomal linker DNA region.** Alignment was performed only on the nucleosome core region. Subsequently, a local mask encompassing both the H1 and linker DNA was applied for 3D classification without alignment, enabling the separation of distinct conformational states in the linker DNA regions.

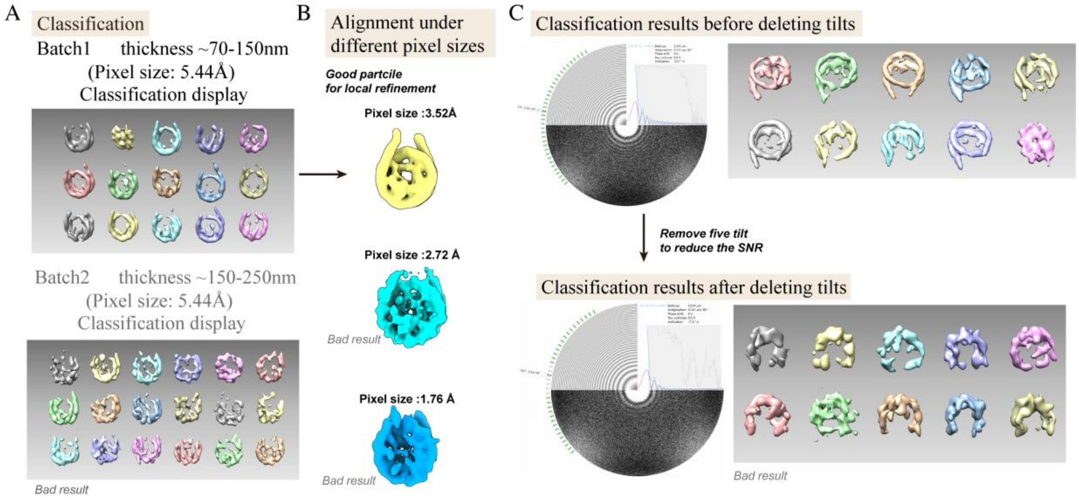


**Figure S4. Impact of SNR on nucleosome classification and alignment quality.**

The lamella thickness, which had a significant influence on the SNR, was crucial to the data quality. Nucleosome feature could be averaged out in the lamella with ~70-150 nm thickness (batch1), but not for those in thicker lamella (~150-250nm, batch2). (A) 3D classification results at different lamella thicknesses. Thinner lamella yield clearer nucleosome features, while thicker lamella result in poorer classification quality. (B) Alignment performance for batch 1 at varying pixel sizes. Higher pixel size (coarser sampling) leads to higher SNR and good alignment; however, reduced pixel size (finer sampling) degrades alignment quality of nucleosomes. (C) Assessment of the influence of SNR on classification. To reduce SNR, five tilt images were removed from the dataset. Classification quality remains high without tilt removal, but deteriorates significantly after removal, indicating that sufficient SNR is critical for accurate nucleosome alignment and classification. The density maps labeled 'Bad result' represent low-quality results.

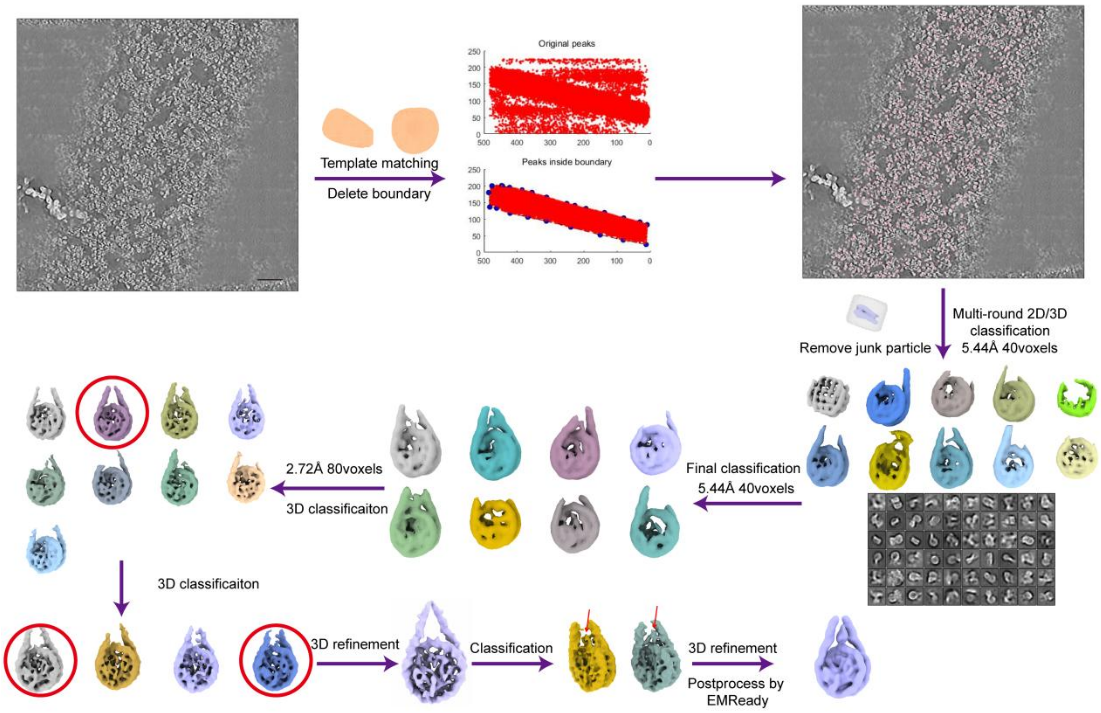


### Figure S5. Flowchart of STA of nucleosomes in lamella.

Nucleosomes within the chromatin in the cryo-FIB lamella are initially selected by template matching. False positive particles are discarded based on the geometrical features of the lamella. The selected nucleosomes are further classified to remove junk particles and improve the overall resolution of the nucleosomes and H1. Scale bars: 50 nm.

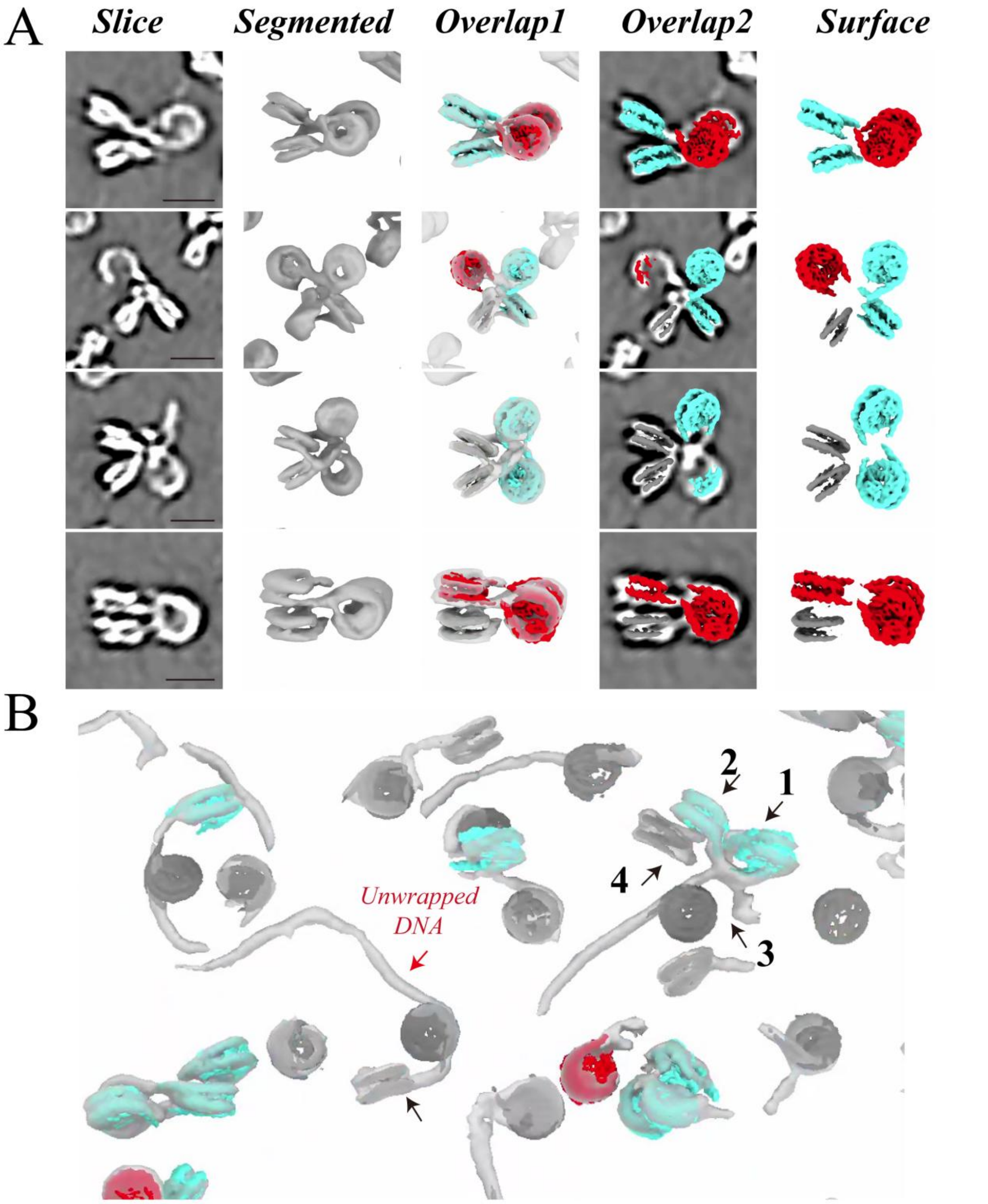


**Figure S6. Visualization of the tetranucleosome structure.**

(A) Visualization of representative tetranucleosome particles. From left to right, each tetranucleosome particle is presented with a tomographic slice, segmented view, an overlay of the segmentation and Nuc-back map, an overlay of the slice and Nuc-back map, and a rotated view for Nuc-back map, respectively. (B) Segmentation of a tetranucleosome tomogram overlaid with the back-mapped nucleosome categories (shown as red, cyan, and black densities) by Nuc-back. A typical unwrapped DNA (red arrow) and an atypical tetranucleosome (black arrows) are shown. Scale bars: 10 nm.

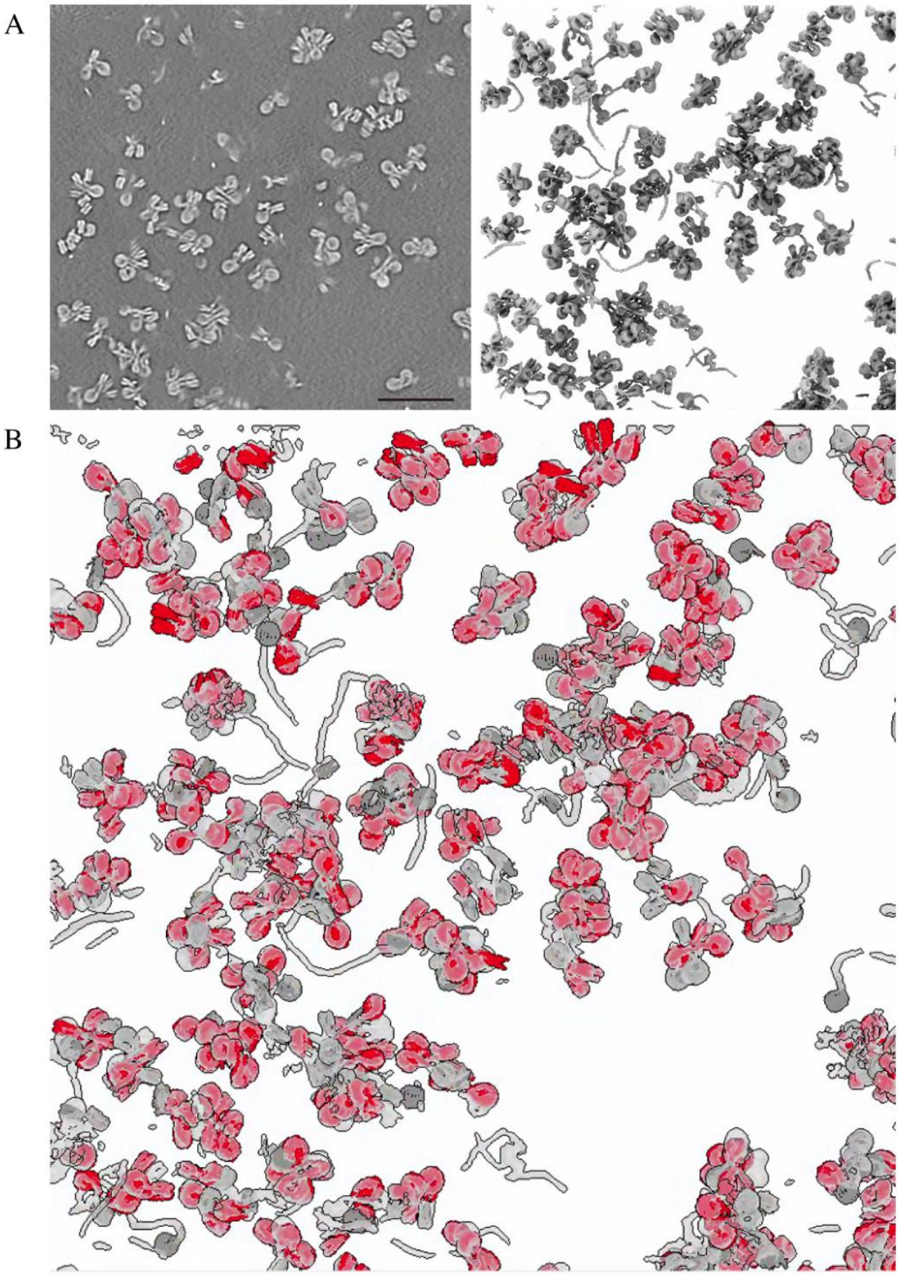

**Figure S7. Visualization of 12× reconstituted chromatin by Nuc-back indicates that most nucleosomes are well located.** (A) From left to right, 12 × reconstituted chromatin is presented with tomographic slice and segmented view. (B) Nucleosomes overlaid on the denoised segment view, showing that the majority of chromatin regions have been successfully occupied by computationally predicted nucleosomes. Scale bars: 50 nm.

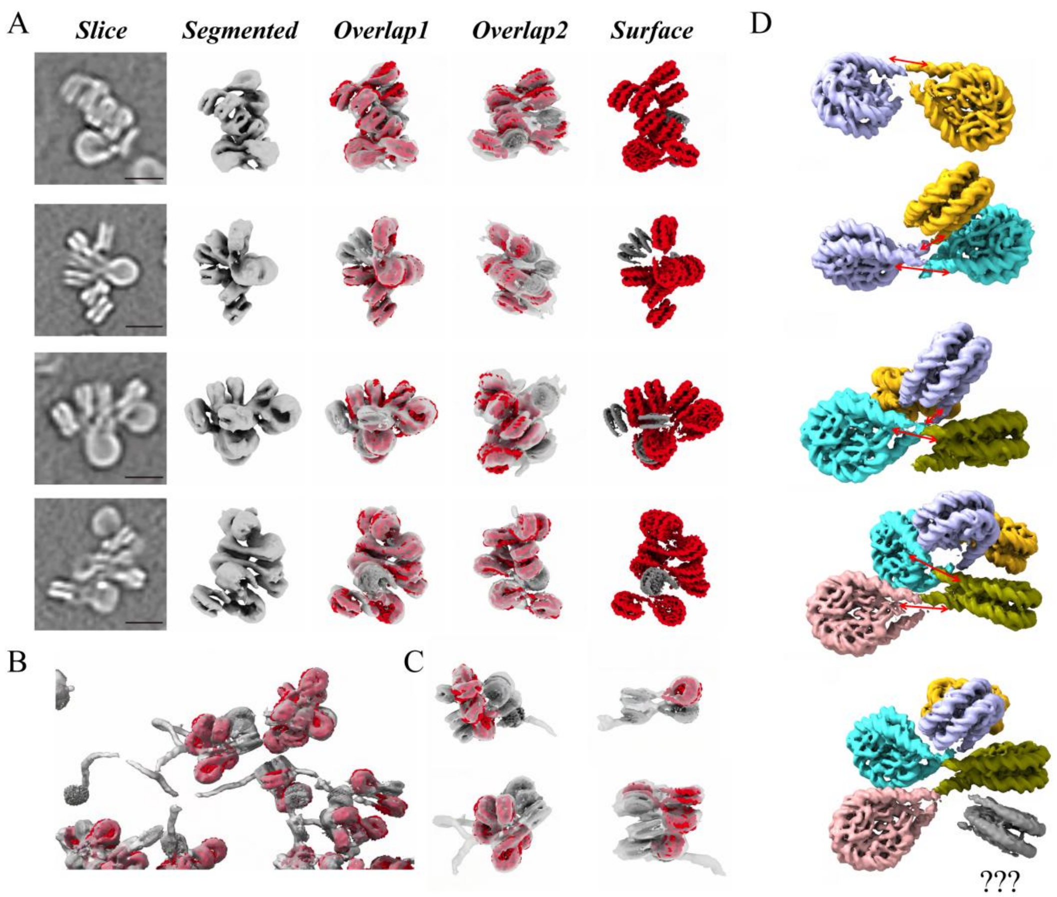


#### Figure S8. Architecture and linker DNA in 12× reconstituted chromatin revealed by Nuc-back.

(A) Visualization of 12× reconstituted chromatin. Each chromatin is presented with a tomographic slice, segmented view, an overlay view of the segmentation and Nuc-back map, a rotated overlay view of the segmentation and Nuc-back map, and the Nuc-back map visualization. (B) A representative tomogram region exhibiting multiple chromatin particles with free DNA unwrapping, which contributes to the structural diversity of chromatin. (C) Representative chromatin particles with unwrapped DNA. (D) The nucleosomes falling with different categories, shown in different colors, are interconnected via averaged linker DNA densities by STA and Nuc-back. The red bidirectional arrows indicate that the 3D density trajectories between nucleosomes can be well traced, but tracing is interrupted at the locations of nucleosomes with poor averaging quality (gray). Scale bars: 10 nm.

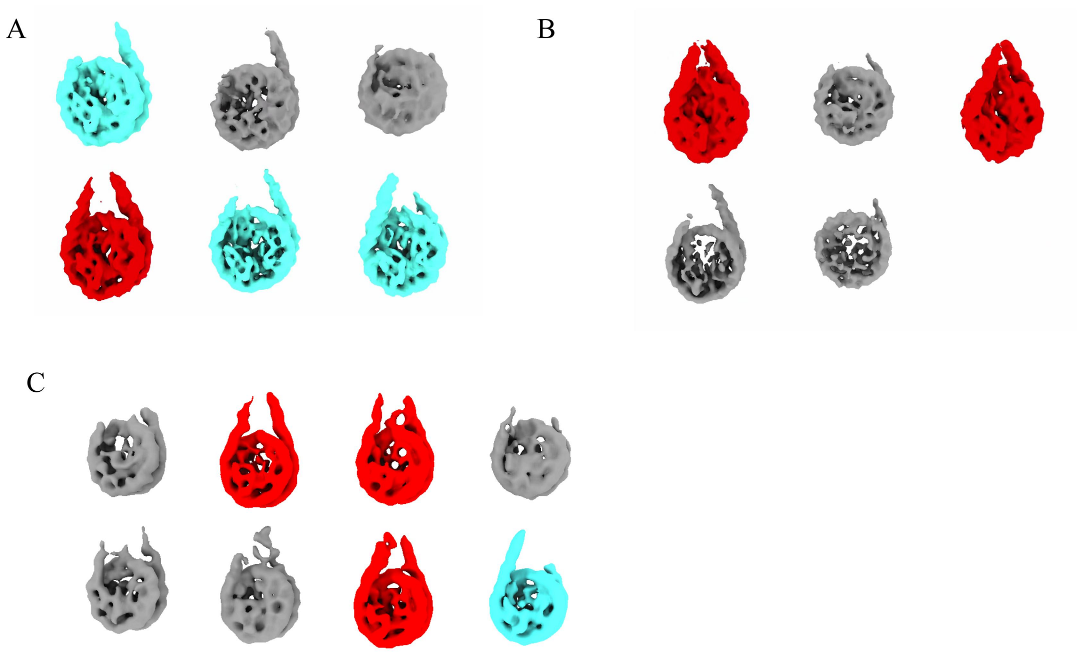


**Figure S9. Classified categories of nucleosome in chromatin from different sources.**

(A) Chromatin extracted from HEK293F cell (dataset 5). (B) Chromatin extracted from HEK293F cell in high salt concentration (dataset 6). (C) Chromatin extracted from HEK293F cell with HAT in a low ratio (dataset 8). The red class corresponds to nucleosomes with linker DNA densities on both sides, the blue class to those with linker DNA on only one side, and the gray class to particles with poorly resolved or missing linker DNA.

**Table S1**

| | 12×177 bp reconstituted chromatin (dataset 1) | Isolated chromatin from SF9 cell (dataset 2) | Chromatin in frog erythrocyte nuclei (dataset 3) | Tetranucleosomes (dataset 4) | Isolated chromatin from HEK293F cell (dataset 5) | Isolated chromatin from HEK293F cell in high salt concentration (dataset 6) | HAT1 with isolated chromatin from HEK293F cell (dataset 7) | HAT2 with isolated chromatin from HEK293F cell in high salt concentration (dataset 8) |
|---|---|---|---|---|---|---|---|---|
| Electron exposure (e/Å$^2$) | ~99 | ~99 | ~99 | ~108 | ~99 | ~99 | ~99 | ~99 |
| Defocus (μm) | -2.8-3.2 | -3.5-4.5 | -3.5-5 | -3.5-4.5 | -3.5-4.5 | -3.5-4.5 | -3.5-4 | -3.5-4 |
| Tilt range (min/max, step) | -50°/+50°, 3° | -50°/+50°, 3° | -60°/+40°, 3° | -55°/+55°, 3° | -50°/+50°, 3° | -50°/+50°, 3° | -50°/+50°, 3° | -50°/+50°, 3° |
| Tomogram used | 108 | 164 | 199 | 152 | 123 | 82 | 162 | 89 |
| Pixel size | 1.36 | 1.36 | 1.76 | 1.76 | 1.76 | 1.76 | 1.76 | 1.76 |
